# Metastatic prognostic ability of lung cancer stromal cells from a single-cell RNA-seq perspective

**DOI:** 10.1101/2023.07.10.548323

**Authors:** Hyunjin Moon, Jonghwan Kim, Hajin Jeon, Jin Ok Yang

**Author notes:** Corresponding author: Jin Ok Yang.

## Abstract

Single-cell RNA sequencing (scRNA-seq) has been widely studied and analyzed to understand cancer heterogeneity. Metastasis and invasion through communication with immune cells have been widely studied; stromal cells are known to change during cancer progression and cause metastasis, but little is known about their inherent metastasis prognostic abilities. This study investigated the abilities of stromal cells by analyzing the scRNA-seq data of adjacent, tumor, and metastasized tissues, biopsied from 15 patients. We considered fibroblast and smooth muscle cells as cell subtypes of stromal cells. We detected decorin (*DCN*) and insulin-like growth factor-binding protein 7 (*IGFBP7*) as sub-cell type markers conserved in tumor and metastasis. We found an organic relationship that affects metastasis by assessing the interaction between the expression and related pathways among the assigned stromal cell subtypes. In addition to the interactions of sub-cell-types within stromal cells, we also studied communication between stromal cells and the five assigned lung cancer cell types, and observed its relation with migration and metastasis; the role of *DCN* as a mediating factor was also studied. Results of our study indicated that *DCN* and *IGFBP7* are factors that can be monitored in the follow-up of prognostic metastasis factors in patients with lung cancer. Therefore, *DCN* and *IGFBP7*, which are the assigned sub-cell types marker in lung cancer stromal cells, can be used as potential biomarkers for follow-up in lung cancer metastasis. Our study assigned stromal cell subtypes of lung cancer and detected markers that can be used as monitoring factors during metastasis of lung cancer. This suggests that *DCN* and *IGFBP7* are potential biomarkers to evaluate metastatic ability and follow-up as metastatic prognostic factors in lung cancer patients.

**Author Summary:** Lung cancer is known to be a disease with high heterogeneity. In particular, it has been studied that stromal cells of lung cancer are involved in metastasis in cancer. Finding specific markers of stromal cell subtypes can lead to more accurate cancer markers. In this study, we assigned stromal cell subtypes as FB and SMC through scRNA-seq data analysis. *DCN* and *IGFBP7*, which are markers specifically expressed in stromal cell sub-cell types of NSCLC and metastatic NSCLC, were found. Whether *DCN* and *IGFBP7* can be used as prognostic factors for metastasis was verified through co-expression analysis of FB and SMC and cell-cell communication. Specific expression as a marker was verified by confirming the role in the hub-gene network and the target genes of communication. Our findings suggest that *DCN* and *IGFBP7* are potential biomarkers for evaluating the metastasis prognostic ability of stromal cells and can be monitored in the follow-up of metastasis prognostic factors in lung cancer patients.

## Introduction

Despite rapid advances in drug development and surgical methods, lung cancer remains the primary cause of cancer-related death worldwide, with an overall 5-year survival rate of approximately 25% [1,2]. Approximately 30% of patients with pathological stage I (p-stage I) non-small-cell lung cancer (NSCLC) develop tumor recurrence and die, despite complete surgical resection [3]. These patients are associated with a poor prognosis because of potential metastasis despite surgical treatment [4].

Moreover, about 40% of patients with NSCLC are diagnosed with NSCLC accounts for 87% of lung cancer cases and approximately 40% of these patients are diagnosed with metastatic disease at presentation, with the most common sites of metastasis being the brain, liver, adrenal glands, and bones [5]. Despite the prevalence of brain metastases, progress in treatment has been slow—hampered by a lack of model systems, a shortage of human tissue samples, and the exclusion of patients with brain metastasis from many clinical trials [6]. Understanding the mechanism of metastasis in lung cancer will help identify new therapeutic strategies.

In lung cancer, the host tissue stroma structurally supports both normal epithelial tissues and their malignant counterparts. Although most host cells in the stroma have specific tumor-suppressing abilities, the stroma undergoes changes during malignancy that facilitate the promotion of growth, invasion, and metastasis in cancer [7]. Cancer-associated fibroblasts (CAFs), the most common type of stromal cells, are an important source of extracellular matrix (ECM)-degrading proteases, such as matrix metalloproteinases (MMPs), which play several important roles in tumorigenesis. Depending on the substrate, MMPs can promote invasion, angiogenesis, recruitment of inflammatory cells, and metastasis through ECM degradation [8]. Accumulating evidence suggests that these stromal components further suggests that these stromal components also regulate drug resistance [9]. Cancer stromal cells have been investigated as potential therapeutic targets because they promote tumor growth through phenotypic changes [10].

The importance of stromal cells in cancer metastasis has been established for numerous cancer types. Approximately 40%–50% of all colorectal cancer (CRC) patients present with metastasis at the time of diagnosis or recurrent disease. In CRC patients, the secretion of IL (Interleukin)-11 from CAF, stimulated by TGF(Transforming growth factor)-β, triggers GP130/STAT3(signal-transducing β-receptor/ transcription factor) signaling. This molecular crosstalk confers a survival advantage to metastatic cells [11]. In hepatocellular carcinoma (HCC), stromal cells have been shown to play an important role in tumor initiation, progression, and metastasis. An increase in angiogenic factors (secreted from tumor cells, macrophages, and activated stellate cells) affects tumor growth by inducing new blood vessel formation [12]. Studies on the role of stromal cells in various carcinomas have revealed that cancer prognosis varies depending on the composition of the microenvironment, and the ratio of stromal cells [13, 14].

Two recent studies in particular have provided detailed gene expression profiling of lung cancer cells. First, data from a study that used single-cell RNA sequencing (scRNA-seq) was combined with multiplex imaging in a large cohort of lung tumors, to show that CAFs participate in T-cell exclusion from tumor nest [15]. Second, data were used from a study in which the loss of developmental stage-specific constraints in macrometastases—triggered by NK cell depletion—was related to the interaction between developmental plasticity and immune-mediated pruning during metastasis. These studies have shown that cancer can induce phenotypic heterogeneity by disrupting tissue repair mechanisms [16].

Despite confirming the importance of stromal cells in lung cancer progression and metastasis, these studies provided limited value in predicting metastasis based on stromal cell profiling [7,9,17–19], because it only identified heterogeneity regarding stromal cell–immune cell communication and phenotype of primary lung adenocarcinoma metastasis. To the best of our knowledge, there have been no studies on the gene expression patterns of lung cancer stromal cell subtypes from scRNA-seq data, the role of different pathways in their expression, the associated gene networks, and the role of stromal cells in communication with other cell types. Therefore, in this study, we utilized these data sets to assess the ability of stromal cells to communicate metastatic potential and to better understand the underlying mechanisms. Our main findings were to assign cell subtypes according to heterogeneity using scRNA-Seq and to evaluate gene expression, cell-cell communication, and co-expression. We understood the mechanisms of the overall metastatic process and detected potential metastasis-mediating markers.

## Results

### Sub-cell type classification of NSCLC stromal cells

To analyze the metastasis prognostic abilities of stromal cells in NSCLC, we performed scRNA-seq analysis of biopsied tissues obtained from 15 patients with lung cancer (Fig 1A) [15, 16]. After clustering in Group 1, in which stromal cells were first selected experimentally from 9 NSCLC patients, only stromal cells were obtained more deeply with 19 clusters through cell type assignment. As a result of the identification of cluster-specific markers, the top 20 markers with the most expression difference for each cluster in Group 1 were detected. The top 20 marker genes in each cluster were used to classify immune and non-immune cells. The immune cells were classified as sub-cell types: granulocyte, lymphocyte, and myeloid cells; non-immune cells were classified as endothelial, epithelial, and stromal cells (Fig 1B). Of these, only the stromal cells were re-clustered based on markers associated with fibroblast (FB) and smooth muscle cell (SMC)-type markers, resulting in the 16 clusters (S1 Table). The sub-cell type of stromal cells was classified into 12 clusters of FB and 4 clusters of SMC (Fig 1C).

**Fig 1.**
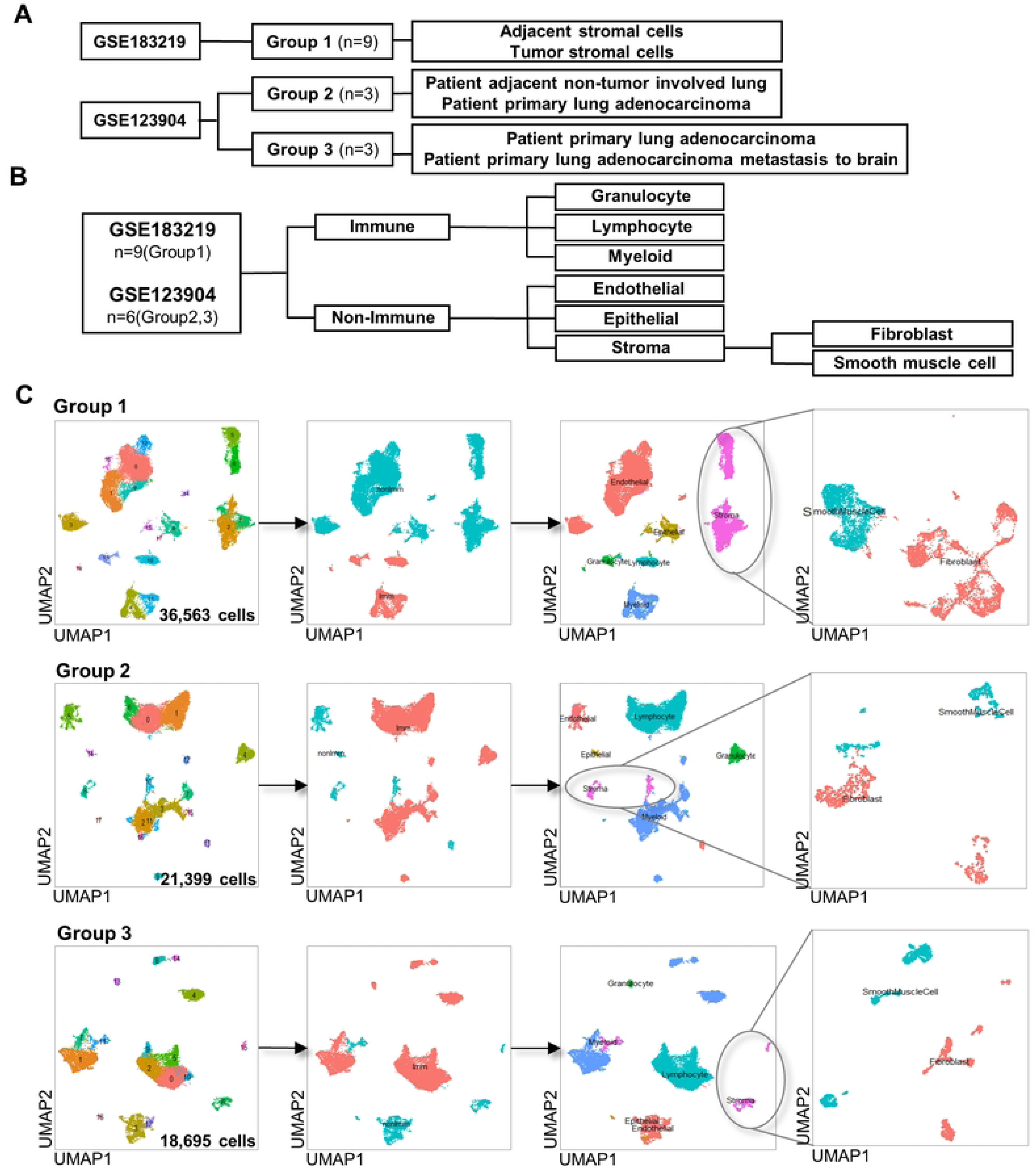
Sample grouping and cell type assignment. (A) Sample grouping (B) Schematic of cell type assignment (C) Clustering result of Group 1 (Ad Stroma vs. Tu Stroma); Group 2 (Ad Non-Tu vs. Pri-Lung Adeno); and Group 3 (Pri-Lung Adeno vs. Pri-Lung Adeno Meta-Brain).

To demonstrate the objectivity of our study, samples were obtained from patients with adjacent non-tumor involved lung (adjacent cell), from patients with primary lung adenocarcinoma (tumor cell) (three samples), and from patients with primary lung adenocarcinoma which metastasized to the brain (metastasis cell) (three samples). Moreover, Group 2 (Ad non-Tu vs. Pri-lung Adeno) and Group 3 (Pri-lung Adeno vs. Pri-lung Adeno meta-brain) were then clustered (Fig 1A).

As a result of clustering in Group 2, 18 clusters were formed. After selecting only stromal cells as the top 20 cluster-specific markers with the most expression difference in Group 2 by cluster, 17 clusters were obtained. Each FB and SMC were classified into nine and eight clusters (Fig 1C).

As a result of clustering in Group 3, 17 clusters were formed. Group 3 was also clustered using only stromal cells as the top 20 cluster-specific markers, and 17 clusters were obtained. FB and SMC were classified with eight and nine clusters, respectively (Fig 1C). As a result of cell type classification, it was observed that decorin (*DCN)* of FB and insulin-like growth factor binding protein 7 (*IGFBP7)* of SMC were the only markers specific to the sub-cell type of stromal cells reproduced in all groups.

### Cell-cell communication

To confirm communication between the cell types, Group 1—in which stromal cells were first selected experimentally—was excluded and cell–cell communication was analyzed [20–22] using Groups 2 and 3. After identifying the top ligands by prioritizing them based on predicted ligand activity, using the Pearson correlation coefficient, it was confirmed which cell types express these top ligands. Based on the percent expressed and log fold change of the expressed ligands, it was confirmed which cell type of ligands communicated with the receptors of the stromal cells and selectively targeted *DCN* and *IGFBP7*.

In Group 2, *DCN*, a marker of FB, communicated with TGF-β1, IL-1B, and IL-1A, which are downregulated ligands in myeloid cells and lymphocytes (Fig 2A), as well as various receptors in stromal cells, thus becoming a target (S2 Table). *IGFBP7* was not targeted by communication between these cell types.

**Fig 2.**
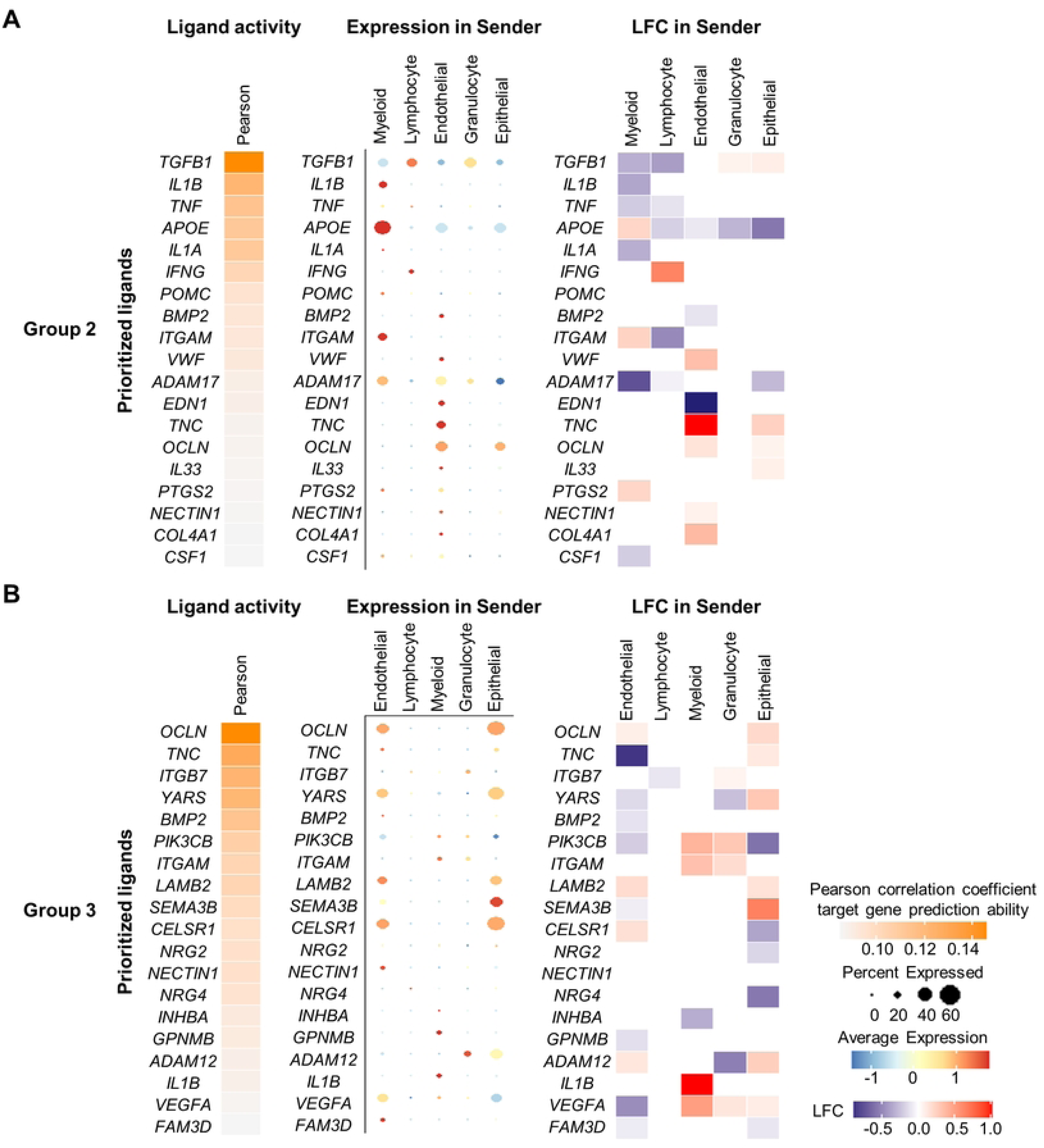
Cell–cell communication results of Groups 2 and 3. (A) In order, top ligands were selected by prioritizing them based on the predicted ligand activity, using the Pearson correlation coefficient, of Groups 2; expression of top ligands by cell type, and expression of ligand according to LogFC in cell types. (B) Top ligands were selected by prioritizing them based on the predicted ligand activity, using the Pearson correlation coefficient, of Groups 3; expression of top ligands by cell type, and expression of ligand according to LogFC in cell types. LogFC; log fold chain.

In Group 3, most of the top ligands were mainly expressed in endothelial, epithelial, and myeloid cells (Fig 2B). *DCN* and *IGFBP7* were not targeted by communication between cell types. The ligands IL1R1 and IL1RAP were the most upregulated in myeloid cells, and IL1B communicated and targeted genes related to metastasis, such as MMP (S2 Table).

### FB and SMC co-expression analysis

For co-expression analysis, differentially expressed genes (DEGs) with similar expression patterns were bundled into the same module using weighted gene correlation network analysis (WGCNA) [23, 24]. Genes that were not grouped by co-expression in stromal cells were excluded, and genes with similar expression patterns were grouped into a module (M). As a result of co-expression of WGCNA in Group 1, 2406 genes formed 12 modules; in Group 2, 4747 genes formed 11 modules; and in Group 3, 3346 genes formed 12 modules (S3-5 Tables).

As a result of the individual module network, the top 10 genes of kME (eigengene-based connectivity), which are hub genes in highly connected modules, formed a hub-gene network and each group’s module had a molecular function corresponding to the characteristics of this hub-gene network. In Group 1, *DCN* and *IGFBP7* belonged to ‘the hub genes of M9 (module 9) in highly connected top 10 genes; in Group 2, *DCN* was not among the top 10 highly connected genes but belonged to M4, whereas *IGFBP7* belonged to the hub genes in highly connected top 10 genes of M2; in Group 3, *DCN* and *IGFBP7* did not belong to the genes composing the module (S3-5 Tables).

The pathway of the module with increased expression was confirmed in FB and SMC of Groups 1–3 (Fig 3A–C). Marker gene overlap analysis was performed using the OverlapModulesDEGs function of WGCNA. The odds ratio (OR) quantifies the correlation between the gene list in the current module and the gene list for the marker gene. In FB and SMC, the related pathways of modules with high OR values were identified, and the pathway which had modules with high OR common in FB and SMC were confirmed (Fig 3D–F; S6-8 Tables).

**Fig 3.**
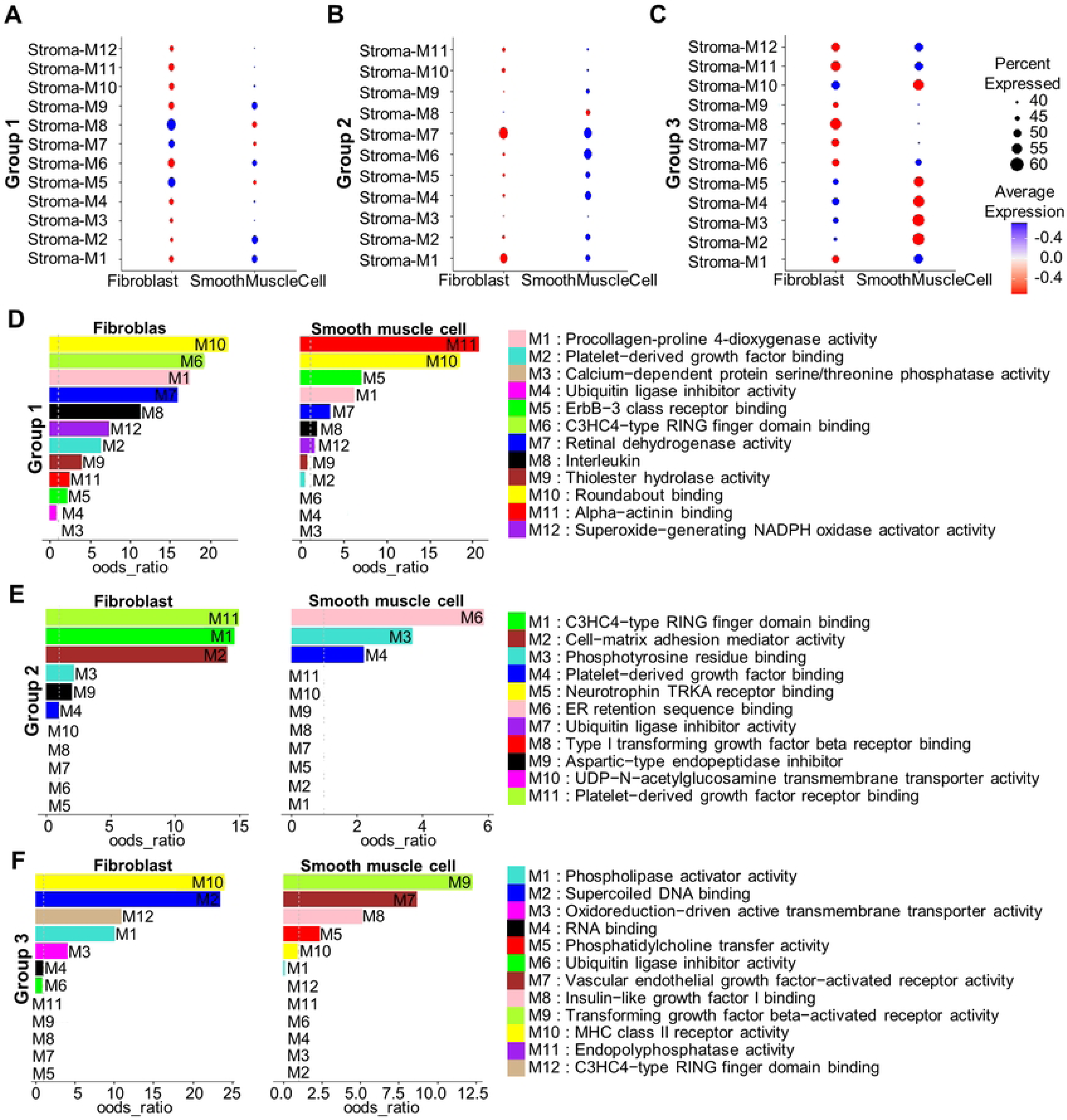
Expression dot plot of FB and SMC according to co-expression analysis and bar plot of FB and SMC according to odds ratio. (A–C) Expression for each module of FB and SMC; red dot indicates an increase in expression, and the blue dot indicates a decrease in expression. (D–F) Results of marker gene overlap analysis for each group. Color bar; GO_Molecular_Function_2021 of each module top-level.

## Discussion

It has been reported that stromal cells of lung cancer affect metastasis [7, 11, 12, 19]. We identified sub-cell type markers that indicate potentially universal characteristics during metastasis in tumor stromal cells, despite the heterogeneity of NSCLC patients in terms of cancer stage and mutation profile. The results of our study provide valuable insights that ease the difficulty of longitudinal analysis of the same patient in cancer studies, using time-series samples. The existing cell type atlas of lung cancer has assigned sub-cell types to two types of smooth muscle cells, five types of fibroblasts, mesothelial cells, and pericytes [25]; in addition, our study identified the cells with functional heterogeneity between the two cell types as FB and SMC. We detected subtype-specific markers of stromal cells through type assignment.

According to early genetic studies on mice, *DCN* deficiency causes angiogenesis inhibitory function in the tumor microenvironment, while overexpression of *DCN* reduces tumor growth rate and suppresses tumor angiogenesis [26, 27]. However, a study [28] on the association between the pathological stage of patients with breast cancer and *DCN* levels in the circulatory system and surrounding tissues reported that *DCN* expression in stromal cells was elevated because stromal and plasma *DCN* levels paradoxically increased in stage II/III patients. Plasma *DCN* can be seen as one of the independent predictors of stage II/III cancer. An increase in circulating *DCN* levels suggests tumor progression; subsequently, proteoglycan synthesis in CAFs shifts from *DCN* to versican, biglycan, and type I collagen, resulting in ECM formation supporting the tumor and allowing it to metastasize. Even if the plasma *DCN* expression is compared with that of the previous studies, plasma *DCN* levels can act as a predictor of progressing disease. In addition, *DCN* was used as a cell-type marker for fibroblast in the advanced NSCLC single-cell atlas and has been demonstrated as a marker significantly expressed in stromal cells in single-cell units of lung cancer [17]. *DCN* expressed in FB, a stromal cell sub-cell type of lung cancer patients, is different from the expression level known in the studies described above. However, scRNA-seq analysis of lung cancer stromal cells suggests that it can be used as a monitoring factor for prognosis and diagnosis of metastasis.

According to previous studies, *IGFBP7*, a cell type marker of SMC, was also observed to be downregulated in various cancers such as breast, gastric, hepatocellular, and colorectal carcinoma [29–32]. In NSCLC array data, the most differentially expressed *IGFBP7* functions as a potential inducer of lymph angiogenesis. Lymphangiogenesis is involved in tissue repair and chronic inflammatory processes as well as tumor lymph node metastasis. Metastatic tumor cells that spread through the lymphatic system are often the most important prognostic factor for patients with carcinoma. Therefore, *IGFBP7* can predict the presence of lymph metastasis in lung adenocarcinoma, aiding in improving the treatment, and can be used as a new tumor marker [33]. Previous studies [26, 27, 29–32] showed downregulation of the expression of *DCN* and *IGFBP7* in tumors rather than in normal cells. However, this was significantly differentially upregulated expressed in patients who underwent advanced cancer therapies, lobectomy, or pneumonectomy for some type of carcinoma. In particular, based on the results of this study showing a significant increase in expression in NSCLC, the possibility that *DCN* and *IGFBP7* can be used as cell type-specific metastasis factors in lung cancer was confirmed. Although it would be challenging to distinguish between patients with and without tumors based on only the expression of *DCN* and *IGFBP7* in lung cancer stromal cells, the expression of *DCN* and *IGFBP7* in the follow-up of such patients with clinically progressing disease may indicate metastasis of cancer.

Through cell–cell communication, we determined which cell types of ligands and stromal cell receptors communicated and targeted *DCN* and *IGFBP7*. In Group 2, *DCN* was expressed as a target for communication with stromal cells, myeloid cells, and lymphocytes, but not in Group 3. This result of cell–cell communications showed that can be used as a monitoring factor *DCN*. Since IGFBP7 was not targeted in all groups, it cannot be considered to be expressed due to communication between cell types. IL-1B, a ligand expressed in Group 3 myeloid cells, is a cytokine that plays various roles in the tumor microenvironment and has been shown to promote metastasis of lung cancer by inducing angiogenesis, tumor epithelial-to-mesenchymal transition, growth, invasion, and adhesion [34]. IL-1B communicates with the stromal cell receptors IL1R1 and IL1RAP and targets key genes that induce metastasis such as MMP and STAT3. The results of this study revealed how communication between cell types occurs in metastasis-induced lung cancer.

Co-expression of WGCNA is the result of overlapping DEGs and marker modules; genes in the same module have similar expression patterns and are involved in pathways [23, 35]. According to the recently known non-canonical function, SRP related to the pathway of the module whose expression increased in FBs of all groups was studied to regulate cell–cell communication, proliferative effect, stress response, and apoptosis regulation [36].

The increased modules in SMC of Group 1 were mainly related to transcription. Abnormally high-level changes in transcription by RNA polymerase II in cancer patients have been used as therapeutic targets [37]. Lack of regulation of ECM components associated with modules with high OR in FB provides cancer progression and metastasis [38]. In SMC, the migratory ability of cells associated with modules with large OR decreased. Stress fiber assembly associated with M10, which has a large OR in FB and SMC, is expressed in FB and plays a central role in cell adhesion [39].

Increased cap-dependent translation due to activation of the mammalian target of rapamycin associated with a module that exhibited increased expression in SMC of Group 2 results in cell proliferation, increased cell survival, and evasion of anti-tumor immunity in cancer cells [40]. The unfolded protein (UPR) associated with a module with increased expression and high OR in FB can play a pro-tumorigenic role by increasing protein folding capacity and prolonging resistance to anticancer drugs [41]. In SMC, neutrophils associated with modules with high OR and reduced expression play a central role in anti-tumor immunity, and the downregulation of neutrophils hinders the normal activity of the immunity system [42].

Microtubules associated with modules with increased expression in SMC of Group 3 contribute to the regulation of cell shape and facilitate the movement of cell organelles and vesicles [43]. The role of protein transport and cellular protein localization associated with a module with reduced expression in SMC and high OR in cancer stromal cells is unknown; hence, further studies are required. The results of WGCNA indicated that expression increased in FB and SMC of groups 1-3 and pathways associated with modules with high OR were all associated with metastasis, migration, and proliferation of lung cancer. Therefore, it was difficult to distinguish the group in which metastasis did and did not occur by pathway alone.

Summarizing the results, the scRNA-seq analysis showed that *DCN* of FB was no longer targeted through cell–cell communication at the time of metastasis. In addition, *DCN* and *IGFBP7* were not grouped as modules in Group 3 where metastasis occurred through co-expression analysis. In Group 3, *DCN* and *IGFBP7* do not belong to the module because they do not form a gene network due to differences in expression patterns from genes significantly expressed in cell subtypes of lung cancer. *DCN* and *IGFBP7* are genes that are significantly expressed in distinguishing cell types in all groups, modules network was not established in tissues where metastasis occurred.

*DCN* and *IGFBP7* were also expressed as stromal cell and SMC markers in brain cancer tissue that metastasized from lung cancer [44]. In normal brain tissue, it was not expressed in the top 30 markers for each cluster [45]. *DCN* and *IGFBP7*, which were identified as stromal cell markers via single-cell RNA analysis, and their role in the mechanism and the mechanism by which lung cancer metastasizes are still poorly understood. However, our findings suggest that *DCN* and *IGFBP7* are potential biomarkers to assess the metastasis prognosis ability of stromal cells, and that can be monitored in the follow-up of metastatic prognostic factors in patients with lung cancer.

## Materials and Methods

### Data source

The Gene expression omnibus (GEO) data repository published GSE183219 (https://www.ncbi.nlm.nih.gov/geo/query/acc.cgi?acc=GSE183219) and GSE123904 (https://www.ncbi.nlm.nih.gov/geo/query/acc.cgi?acc=GSE123904) that were used for analysis. GSE183219 provides the biopsy data of nine samples of adjacent stromal cells and nine samples of tumor stromal cells of patients with NSCLC. These samples were collected and immune cells and epithelial/tumor cells with CD45 and Epithelial cell adhesion molecule (EpCAM) were removed. Thus, stromal cells with CD29 and PDPN were positively selected. The data were then sorted. For GSE123904 data, three adjacent tissue samples (Stage IA, IB, IV), three tumor tissue samples (Stage IA, IB, IV, and have mutation EGFR, KRAS), and three patient primary lung adenocarcinoma metastasis to brain tissue(All sample stage IV, and have mutation EGFR, TP53) sample data were obtained from patients with NSCLC.

### Data acquisition and pre-processing

Data processing and analysis of scRNA-seq data were performed using R (v.4.1.3) [46] Seurat packages. Unless otherwise specified, data from all groups were analyzed using the same options. Low-quality/dying cells often show contamination of mitochondria, so quality control was performed to remove low-quality events (total number of RNA > 200, percent of mitochondria < 20). To normalize the data set, LogNormalize, a global-scaling normalization method, was used that normalizes the feature expression measurements of each cell to the total expression, multiplies the scale factor (10,000 by default), and converts the result to log. To reduce the batch between samples, the batch collection was performed using harmony packages. Cluster markers were identified using the FindAllMarkers function (MAST test, with default settings treating nUMI and ambient RNA content as latent variables).

Cluster markers were identified using the FindAllMarkers function (MAST test, with default settings treating nUMI and ambient RNA content as latent variables). Stromal cells were identified within the cluster using all of the top 20 markers with the most expression difference based on Log2FC.

### Gene ontology and pathway enrichment analysis

Gene Ontology (GO) enrichment analysis of hub-genes co-expressed for each module was performed using the R package enrichR (v.3.1) GO_Biological_Process_2021, GeneOverlap (v. 1.30.0) [23, 24] and GetEnrichrTable function was used for enrichr pathway enrichment analysis. The GO consists of three components: biological process (BP), molecular function (MF), and cellular component (CC). The pathways of overlapping module genes in FB and SMC were defined by Enrichr Ontologies’ GO Biological Process 2021.

### Weighted gene co-expression network analysis (WGCNA)

To construct a co-expression network of genes and detect modules, a Seurat object was set up using the SetupForWGCNA function [23, 24], and a gene was selected for use. For the fraction option, genes expressed in specific cells in the entire dataset or each cell group were used. Subsequently, the Metacells were constructed from a single-cell dataset. The Metacells are aggregates of small groups of similar cells derived from biological samples of the same origin. The Nearest Neighbors (KNN) algorithm was used to identify similar cell groups and calculate the average and sum of cell expressions to create a Metacells gene expression matrix, orig.ident (Group 1-3), with cells grouped by type.

We selected a mild threshold to construct the normalized meta-cell expression matrix. WGCNA was used to prepare a gene-gene correlation adjacency matrix to infer co-expression relationships between genes. It is important to designate an appropriate, mild threshold to reduce the amount of noise in the correlation. The best power value of each group was determined as adjacent stromal cell-tumor stromal cell 7, adjacent cell-tumor cell group 10, and tumor cell-metastasis cell 20. Based on this, a scale-free network and a topology overlap matrix (TOM) were constructed. Since the ‘gray’ color is composed of genes not grouped into co-expression modules, grey-colored modules were excluded from all downstream analyses and interpretations. Module Eigengenes (MEs) is a commonly used metric to summarize the gene expression profile of the entire co-expression module. This process was performed to reduce technical artifacts generated during dimensionality reduction.

The ModuleNetworkPlot function was used to visualize the network underlying each module’s top 25 hub genes. In this network, each node represents a gene, and each edge represents a co-expression relationship between two genes in the network. The top 10 hub genes of kME were placed in the center of the plot, and the remaining 15 genes were placed in the outer circle. In the marker gene overlap analysis process, we compared the WGCNA module with the cluster or cell-type marker gene. First, Seurat’s FindAllMarkers was used to identify marker genes for each cell type, and then overlapped modules and DEGs were plotted using OverlapModulesDEGs. Each point of overlap of modules and cell-type markers is designated with a unique R packages hdWGCNA module color; the points’ size was adjusted according to the overlap statistic. FDR significance levels are indicated by asterisks above the dots. ‘***’: 0–0.001, ‘**’: 0.001–0.01, ‘*’: 0.01–0.05, ‘+’: 0.05–0.1, (No symbol): 0.1–1.0.

## Acknowledgments

This research was supported by the Basic Science Research Program of the National Research Foundation of Korea (NRF), funded by the Ministry of Science and ICT (NRF-2020M3E5D708517212 and 2020M3A9I6A0103605713).

## Supporting Information

**S1 Table. FB and SMC cell type markers.** Cell type markers of Groups 1-3.

**S2 Table. Cell-cell communication-associated genes.** Stromal cell receptors of ligands TGFB1, IL1B, and IL1A in Group 2, IL1B-IL1R1, and IL1B-IL1RAP communication target genes in Group 3.

**S3 Table. The number of genes per module in Group 1.** Hub genes in highly connected top10 genes and GO_molecular_function.

**S4 Table. The number of genes per module in Group 2.** Hub genes in highly connected top10 genes and GO_molecular_function.

**S5 Table. The number of genes per module in Group 3.** Hub genes in highly connected top10 genes and GO_molecular_function.

**S6 Table. Co-expression in Group 1.** Modules with increased expression in FB, modules with increased expression in SMC, modules with high OR in FB, modules with high OR in SMC, and modules with high OR in both FB and SMC.

**S7 Table. Co-expression in Group 2.** Modules with increased expression in FB, modules with increased expression in SMC, modules with high OR in FB, modules with high OR in SMC, and modules with high OR in both FB and SMC.

**S8 Table. Co-expression in Group 3.** Modules with increased expression in FB, modules with increased expression in SMC, modules with high OR in FB, modules with high OR in SMC, and modules with high OR in both FB and SMC.

